# Using animal tracking for early detection of mass poisoning events

**DOI:** 10.1101/2024.11.29.625987

**Authors:** T. Curk, A. Santangeli, W. Rast, R. Portas, G. Shatumbu, C. Cloete, P. Beytell, O. Aschenborn, J. Melzheimer

**Affiliations:** Leibniz Institute for Zoo and Wildlife Research, Department of Evolutionary Ecology, Berlin, Germany; Animal Demography and Ecology Unit, Institute for Mediterranean Studies (IMEDEA), CSIC-UIB, 07190, Esporles, Spain; FitzPatrick Institute of African Ornithology, University of Cape Town, Cape Town, South Africa; Ministry of Environment, Forestry and Tourism, Directorate of Parks and Wildlife, Subdivision (Airwing), Etosha National Park, Namibia; Ministry of Environment, Forestry and Tourism, Etosha Ecological Institute, Etosha National Park, Namibia; Ministry of Environment, Forestry and Tourism, Directorate of Scientific Services, Etosha National Park, Namibia

**Keywords:** human-wildlife conflict, local enhancement, sentinel animals, sentinel poisoning, social foraging, Gyps vultures, agent-based model

## Abstract

1 Amidst the sixth mass extinction, some groups, such as vultures, the only obligate scavengers among vertebrates, are disappearing at an unprecedented rate. Vulture populations worldwide are largely threatened by poisoning. As many vulture species are social foragers, they can congregate in large numbers to scavenge at a carcass, potentially increasing their exposure to poisoning risk. Current anti-poisoning prevention and mitigation measures are insufficient to tackle this threat. There is an urgent need for new effective strategies to prevent mass vulture mortality.

2 In this study, we used agent-based modeling to: (1) quantify the impact of different foraging strategies on vulture poisoning risk, and (2) evaluate the cost-effectiveness of using vultures as sentinels for poisoning detection. This approach involves GPS tracking of various numbers of vultures and using the data to quickly detect poisoning incidents and decontaminate carcasses. These actions help mitigate further vulture mortality and prevent mass poisoning.

3 Our results indicate that social foraging significantly elevates the risk of vultures being poisoned. Vulture tracking contributes substantially to the early detection of poisoning events, which results in reduced mortality. Poisoning can be mitigated by tracking more vultures and responding faster, with an optimal cost-effectiveness achieved by tracking 5% of the vulture population (25 individuals in our system) and a budget of 58,728 USD. This approach could potentially prevent 45% of poisoning-related mortalities if interventions occur within 2 hours.

4 *Synthesis and applications*: Our results suggest that, in order to reduce mortality incidences from poisoning, it is sufficient to track a small proportion of the vulture population, which would act as sentinels for the rest. By evaluating the costs and ecological benefits of these mitigation strategies, we provide evidence-based solutions that practitioners can use to design conservation plans. These findings are therefore instrumental in supporting vulture and scavenger conservation policy and practice.

## Introduction

Global vulture populations declined drastically in the last decades primarily due to poisoning (Ogada et. al., 2012). The most common poisons used in Africa are agricultural pesticides such as organophosphates and carbamates, which kill vultures almost immediately after consumption (Ogada et al., 2014; 2016a). There are different reasons and motivations for poisoning.

Intentional poisoning includes actions for belief-based use, the bushmeat trade, or the elimination of vultures’ sentinel role in revealing poaching locations i.e., sentinel poisoning (Botha et al., 2017). Unintentional poisoning occurs due to human-wildlife conflict (HWC), where farmers unintentionally poison vultures while targeting predators that killed their livestock. (Ogada, 2014; Santangeli et al., 2016, 2017). While the two types of poisoning, intentional and unintentional have different drivers, they can result in a similar outcome – mass mortality of vultures where dozens or even hundreds of vultures can die during a single poisoning event (Murn & Botha, 2018). A number of such incidences were reported by the African Wildlife Poisoning Database (The Endangered Wildlife Trust and the Peregrine Fund, 2023), including a mass poisoning event in Guinea-Bissau (Henriques et al., 2020).

Current anti-poisoning measures, such as surveillance for prevention, and rapid decontamination of carcasses for mitigation of mortalities, are insufficient to tackle poisoning and prevent population declines (Ogada et al., 2016a; Murn & Botha, 2018; Henriques et al., 2020). Mitigating mortalities when poisoning occurred entails a very rapid detection of poisoning incidences (Murn & Botha, 2018). This allows fast decontamination of the location, preventing further vulture, and other wildlife, poisoning from carcasses that could otherwise last hours to days. While poisoning detection might be efficient in small protected areas with rangers regularly patrolling the site, it is more difficult in large remote areas and extensive ranchland. Here, patrolling is challenging due to terrain, time constraints, lack of work force and resources. In such challenging environments, technology development such as drones, acoustic monitoring, and sentinel animals bearing a GPS tag could greatly help (Mulero-Pázmány, 2014; O’Donoghue & Rutz, 2016; Astaras et al., 2020; de Knegt et al., 2021; Mesquita et al., 2023). Vulture tracking proved to be efficient at detecting herbivore carcasses and identifying associated illegal activities such as illegal carcass disposal in the field and potentially also poaching and poisoning (Mateo-Tomás et al., 2023; Peters et al., 2023; Rast et al., 2024). The use of such new technologies could serve as early warning systems for local authorities. However, their feasibility and cost-effectiveness are still largely unquantified.

Vultures, as obligate scavengers that feed on carcasses (Mundy et al., 1992), are known to utilize social information to locate food. Studies have shown that vultures detect carcasses by following informed individuals from communal roosts, foraging in networks, or approaching individuals that have detected the carcass from kilometers away (Rabenold, 1987; Deygout et al., 2010; Cortés-Avizanda et al., 2012). Three hypotheses have been suggested as possible foraging strategies in vultures (Wittenberger & Hunt 1985; Houston 1974; Jackson, Ruxton & Houston, 2008; Cortes-Avizanda et al., 2014): (1) nonsocial, where vultures use only individually acquired information to find carcasses; (2) local enhancement, where, in addition to individually acquired information, vultures also use social information by approaching other individuals feeding at the carcass or those landing at the carcass; and (3) chains of vultures, where in addition to using individually acquired information and local enhancement, vultures form chains as each individual follows another toward the same food source from kilometers away. Testing the impact of these foraging strategies on poisoning risk and assessing the cost-effectiveness of sentinel vulture-based mitigation is crucial and timely. This has not been addressed before, leaving us to design conservation strategies with limited information. Quantifying cost-effectiveness is essential, especially given the chronic scarcity of conservation resources in the Global South (Obura, 2023; Botha et al., 2024). Such information is also key to securing financial support by demonstrating positive returns to potential funders (Santangeli & Sutherland, 2017).

Here we contribute timely and relevant evidence to fill the above knowledge gaps by quantifying the exposure to a specific threat, and the cost-effectiveness of animal tracking to mitigate this threat, using vultures and poisoning as an example. Specifically, we used agent-based modelling integrated with vulture tracking to (1) assess how different foraging strategies i.e. nonsocial, local enhancement and chains of vultures affect poisoning risk in vultures, and (2) evaluate the cost-effectiveness of vulture tracking for early detection of poisoning events. In doing so, we tested alternative scenarios based on varying poisoning incidence rates, proportion of vultures being GPS tracked, and response time from poisoning detection to decontamination.

## Materials and methods

### Species and study area

We used the white-backed vulture (*Gyps africanus*) as our model organism due to its gregarious feeding habits (Mundy et al. 1992). This species is classified as Critically Endangered by the IUCN, with rapidly declining populations (BirdLife International 2022). White-backed vultures are found across Africa, particularly in Etosha National Park (ENP) in northern Namibia, the focal area of this study. ENP spans 22,270 km² and is home to diverse wildlife while the Etosha Pan is a barren area of 4,730 km² within the park which lacks wildlife. The presence of iconic species attracts poachers to the park, increasing the risk of sentinel poisoning. Additionally, predators from the park often spread to surrounding commercial livestock farmland, heightening the risk of unintentional vulture poisoning due to human-wildlife conflict (Santangeli et al., 2016).

### Vulture tracking and carcass detection

Between 2017 and 2022, we equipped 30 white-backed vultures with Bird Solar GPRS/UMTS 42g units (e-obs GmbH). Most of the tags (n = 27) collected GPS data every minute for 7 seconds at 1Hz, and also ACC (acceleration) data at 20Hz for 6 seconds. The rest of the tags (n = 3) collected GPS data every minute one position and ACC data at 20Hz for 4 seconds. Data were collected only during the day (between 6:00 and 20:00 local time) since vultures are not active at night. After data cleaning, the dataset included 30 individuals between May 1, 2022 and June 1, 2023.

To locate carcasses in ENP, we utilized several machine learning algorithms on vulture GPS and ACC data. Initially, we inferred vulture behavior from ACC data using a Support Vector Machine (Cortes and Vapnik 1995) to classify six behaviors: feeding, grooming, lying, standing, active flight, and passive flight. We validated this by tagging two vultures at Tierpark Berlin and recording their behaviors with cameras. Using BORIS software (Friard & Gamba 2016), we labeled ACC data from the two individuals based on video observations, resulting in a training dataset of 14,682 samples. Comparison with predicted behaviors showed high classification accuracy (0.86 – 1). Next, we applied the DBSCAN clustering algorithm (Ester et al. 1996) to link classified behaviors to GPS positions, identifying clusters with specific behaviors. Each cluster’s center coordinates and behavior proportions were determined. Finally, we used the Extreme Gradient Boosting algorithm (Chen & Guestrin 2016) to differentiate between feeding and non-feeding clusters. Field verification of 1,864 feeding clusters confirmed feeding evidence at 548 clusters, with an overall precision of 93%. The algorithm identified 2,205 feeding clusters in ENP from May 1, 2022, to June 1, 2023, which we used as carcass locations for further analysis. For the updated version of the method, see Rast et al. (2024).

### Agent-based model

We modified and extended the agent-based model in Curk et al. (2025) which simulates foraging and carcass detection of white-backed vultures in ENP. This model is based on Cortés-Avizanda et al. (2014) where the three submodels represent different foraging strategies, nonsocial, local enhancement and chains of vultures (Fig. 1). The model was built in Python and is described in detail in the Overview, Design concepts and Details (ODD) protocol in the Suppl. mat. (Grimm et al. 2006, 2010, 2020, Railsback & Grimm 2019). To parameterize the model, we used vulture tracking data, field observations and literature (Table 1, Suppl. material). We also performed local sensitivity analysis (for details see Suppl. material).

**Figure 1:**
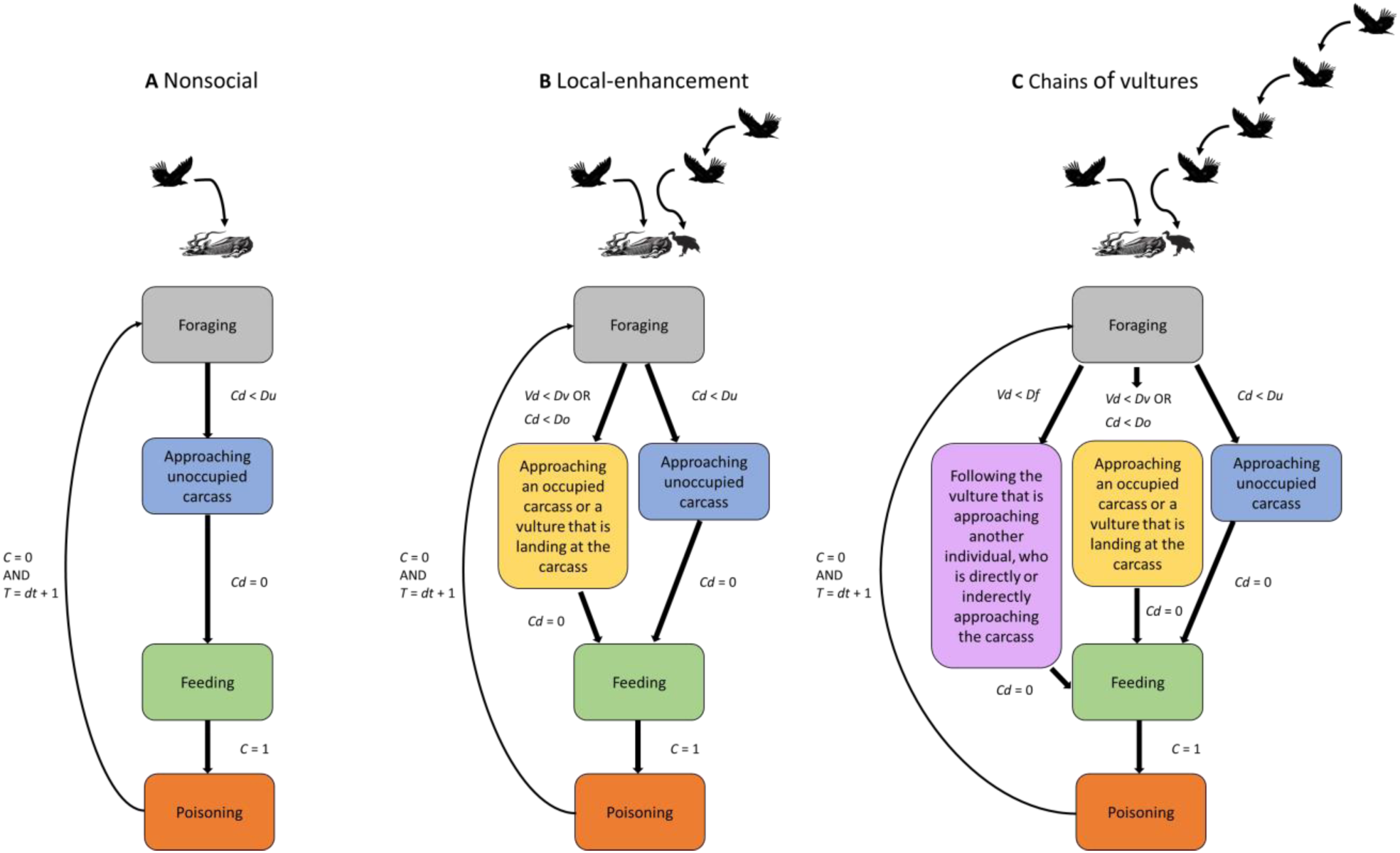
Changes in vulture status in the three submodels adapted from Curk et al. (2025). *Cd* is distance to the occupied or unoccupied carcass and *Vd* distance to a vulture who is landing at the carcass or a vulture who is approaching another individual who is directly or indirectly approaching the carcass. *C* is carcass status (0 – not poisoned, 1 – poisoned), *T* simulation time and *dt*, current date. *Du*, *Do*, *Dv*, and *Df* represent 300 m, 4,000 m, 2,000 m and 2,000 m, respectively.

**Table 1:**
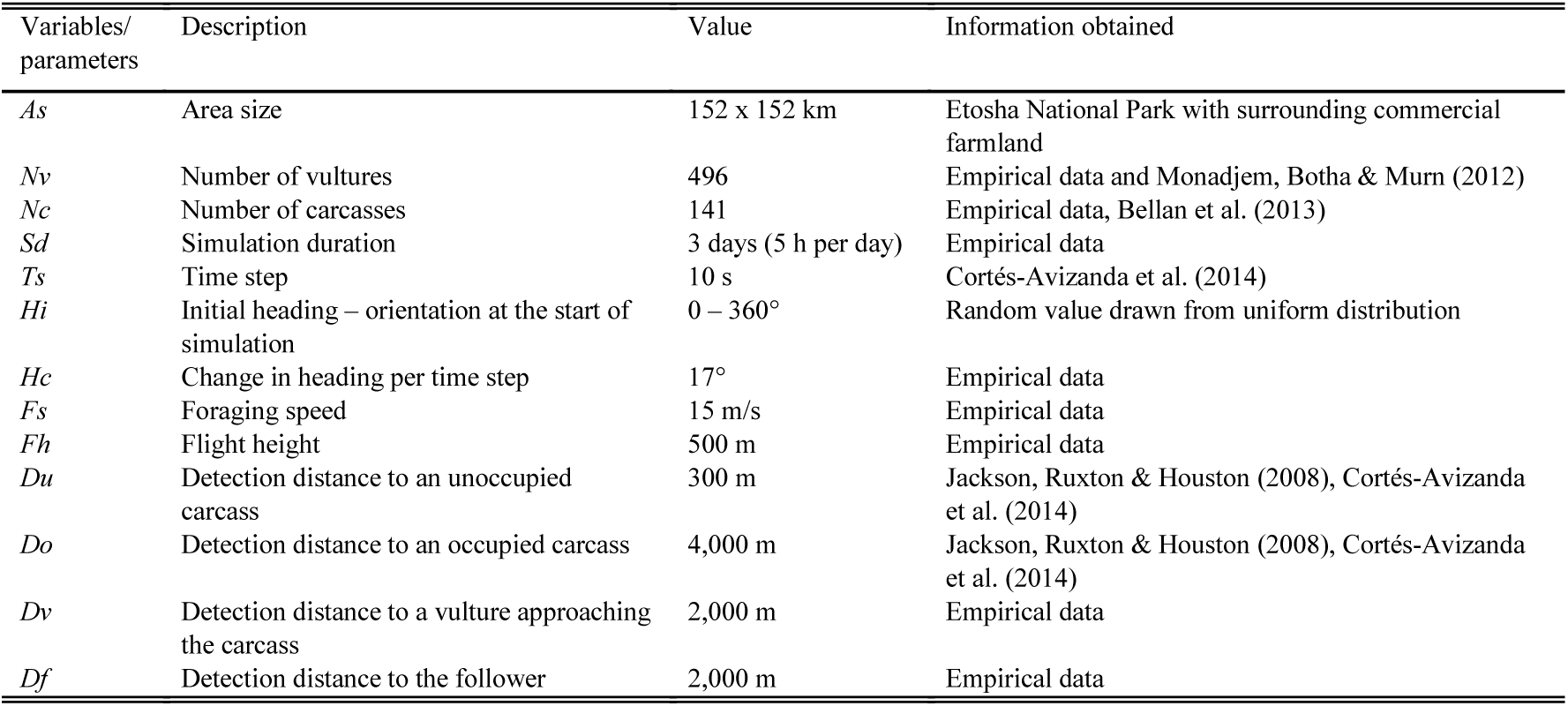
State variables and parameters (adapted from Curk et al., 2025)

The model simulates white-backed vultures foraging over a 152 x 152 km area, representing ENP (excluding Etosha Pan) and a 10 km belt of commercial farmland surrounding the park, over three foraging days, each lasting 5 hours (*Sd*). Carcasses and vultures are randomly positioned in a simulation space. At the start of each simulation, carcasses are randomly categorized between poisoned and not poisoned and vultures as tagged or not tagged. The model operates in discrete 10-second time steps (*Ts*), updating vulture locations at each step. Vultures forage at a constant height (*Fh*), speed (*Fs*), and heading change (*Hc*). Vulture status is initially: 0 – foraging. If the focal vulture is within detection distance from a carcass (*Do, Du*), from another vulture (*Dv*), or from a follower (*Df*), the focal vulture changes direction and approaches the target, changing carcass status from 0 – unoccupied to 1 – detected, and the vulture status to 1 – approaching an unoccupied carcass, 2 – approaching an occupied carcass or an individual who is landing at the carcass, or 3 – following the vulture that is approaching another individual who is directly or indirectly approaching the carcass. The "nonsocial" submodel includes only *Du*, "local enhancement" includes *Du*, *Do*, and *Dv*, and "chains of vultures" includes all detection distances. Vultures are assumed to lose altitude while approaching the carcass from detection distance, so no extra steps for descent are included. When the vulture reaches the carcass, the carcass status changes to 2 – occupied and vulture status to 4 – feeding. If the carcass is poisoned, the vulture status changes to 5 – poisoning and the vulture stays at the carcass until the end of simulation. If the carcass is not poisoned, the vulture status changes next day to 0 – foraging and the states can change repeatedly as explained above.

Using the agent-based model, we first examined how three foraging strategies (nonsocial, local enhancement, chains of vultures) affect poisoning risk in vultures. We ran each submodel multiple times across various percentages of poisoned carcasses (1%, 5%, 10%, 25%, 50%, 100%), totaling 180 simulations (3 submodels x 6 scenarios x 10 repetitions). Next, to assess the effectiveness of vulture tracking for early poisoning detection, we focused on the chains of vultures submodel with varying percentages of poisoned carcasses and tracked individuals, resulting in 360 simulations (1 submodel x 6 carcass scenarios x 6 tracked vulture scenarios x 10 repetitions). We selected the chains of vultures submodel because it incorporates both nonsocial and social strategies, as Curk et al. (2025) indicated that white-backed vultures in ENP have the potential to use different strategies depending on carcass density and availability. This means that if density of foraging individuals is high enough and carcass availability is low, vulture chains can form while if density of foragers is low, the nonsocial strategy is used.

### Model output

To evaluate how different foraging strategies (nonsocial, local enhancement, and chains of vultures) affect poisoning risk, we first calculated the proportion of vultures feeding within the first day for each submodel, considering those with statuses 4 (feeding) or 5 (poisoning) relative to the total number of vultures in the simulation. Next, we quantified the proportion of poisoned vultures at the simulation’s end by assessing the number with status 5 (poisoning) against the total number of vultures.

To assess if vulture tracking enables early detection of poisoning, we calculated the proportion of saved vultures under various response times (immediate, 15 min, 30 min, 1 hr, 2 hrs, 6 hrs, 12 hrs, 1 day, and 2 days). Response time 0 indicates when the first tracked vulture arrives at the carcass and response time 15 min is 15 min after the first vulture arrives at the carcass etc. A carcass is considered decontaminated once the specified response time begins, meaning vultures arriving after this point are considered ’saved’ from poisoning.

We compared model outputs with empirical data, calculating the average number of individuals feeding per day from the carcass detection dataset (mean = 14.5, SE = 0.2), representing half of the 30 tracked vultures. We estimated that this percentage reflects the entire population, suggesting 248 out of 496 vultures feed daily. Initially, from May 1, 2022, to June 1, 2023, we tracked 30 vultures (6% of the population) with one poisoning case (3%). By June 1, 2024, 79 vultures were tracked (16% of the population) with three poisonings (4%). We assumed the same poisoning percentages apply to the entire population, estimating 15 and 20 poisoned vultures, respectively.

### Cost-effectiveness

We quantified the marginal cost-effectiveness of each scenario, assessing how saved vultures and associated costs changed with increasing proportions of tracked vultures. For each response time, we calculated the mean differences in vultures saved and costs between successive tracking proportions across different simulation runs. Marginal cost-effectiveness was determined by dividing the change in saved vultures by the change in costs, indicating how many additional vultures were saved per dollar spent. Costs included Bird Solar GPRS/UMTS units, cellular access, data transmission, harness materials, technician salaries, and fuel. For cost estimation, we followed Iacona et al. (2018) (see “Costs of vulture tracking,” including Table S2 in the Supplement).

## Results

The results of the simulations indicated that nearly all vultures were feeding at the end of the 1-day simulation period in the two social submodels (chains of vultures and local enhancement). In contrast, about 60% of vultures were feeding in the nonsocial submodel (Fig. 2a). This finding aligns closely with our empirical estimate (see “Model output”), which suggests that only half of the vulture population feeds daily.

**Figure 2:**
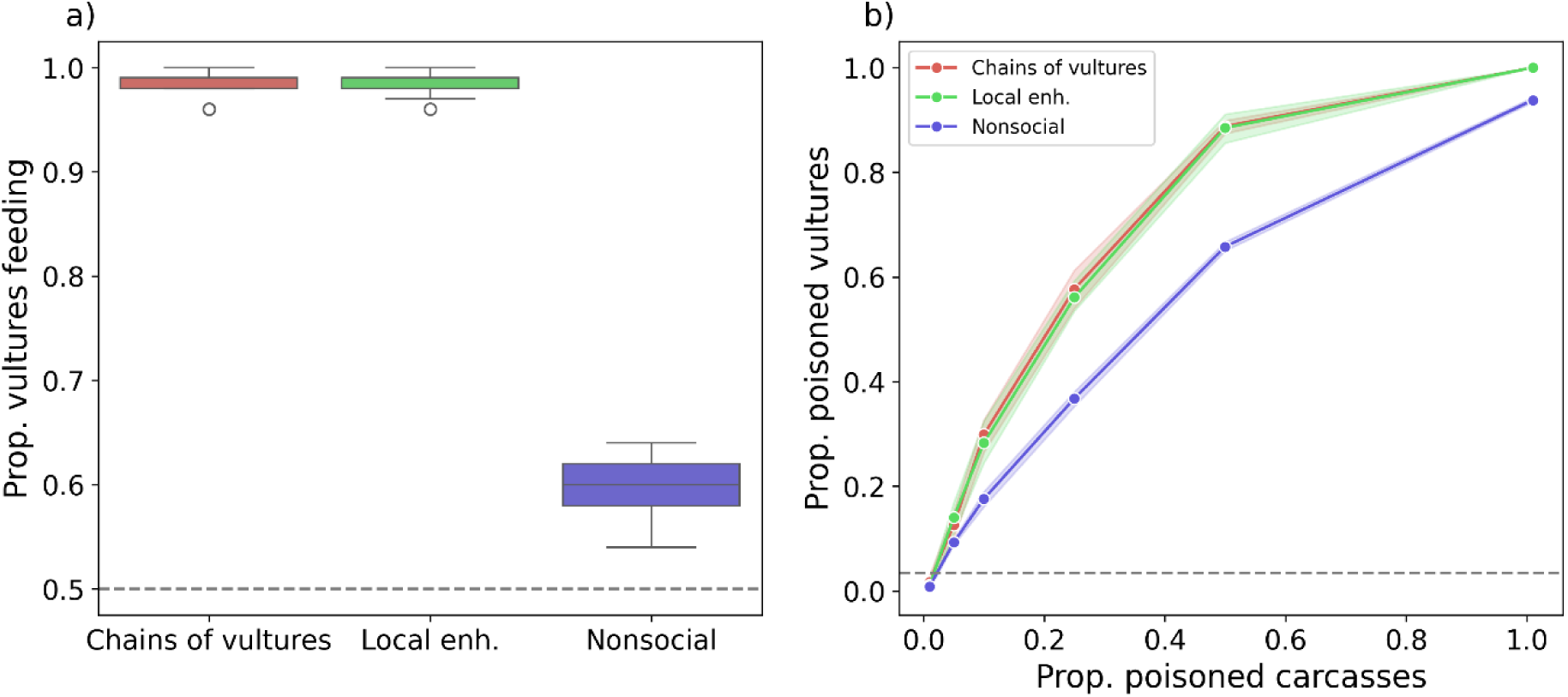
(a) Proportion of vultures feeding after one-day simulation for the three submodels i.e. foraging strategies. (b) Proportion of poisoned vultures under different scenarios of the proportion of poisoned carcasses for the three submodels. Dashed lines represent empirical estimates of (a) the proportion of vultures feeding on average per day in our study area and (b) the proportion of poisoned vultures.

The proportion of poisoned vultures varied based on the proportion of poisoned carcasses and the foraging strategy. Higher poisoning, and social foraging (chains of vultures and local enhancement), strongly elevated the proportion of poisoned birds in the landscape (Fig. 2b). Our estimated proportion of poisoned vultures in the study area was very low (0.03 – 0.04), overlapping with all submodels at the 0.01 proportion of carcasses.

The proportion of tracked vultures in the modeled population played a significant role in preventing poisoning-related mortalities, by early carcass detection and decontamination that can save the lives of incoming birds (Fig. 3). As expected, higher proportion of tracked vultures and lower proportion of poisoned carcasses in the environment resulted in higher proportion of prevented mortalities (Fig. 3a). However, we also showed that to increase the number of saved individuals and prevent large vulture mortalities, the response time to access and decontaminate the poisoned carcass is critical (Fig. 3b). The faster the response time, the more vulture deaths are prevented. Most relevant, the increase in the proportion of saved vultures with the increased proportion of tracked vultures, and the faster response time, was not linear, but asymptotic. This indicates that even small conservation efforts (tracking a small portion of the population, e.g. from 1 to 5% and reducing the response time from two days to < 12 hours) could yield disproportionately high returns (in terms of vultures saved from poisoning) in relation to the costs. Within our estimated proportion of tracked vultures (6 to 16%) in the study area, the model suggests that up to 65% of the vulture population can be saved from poisoning, depending on the response time. For example, if response time to the initial poisoning detection is within 1 hour, more than 50% of the vulture population could be saved from poisoning. If response time is 12 hours, 25% of the vulture population could be saved.

**Figure 3:**
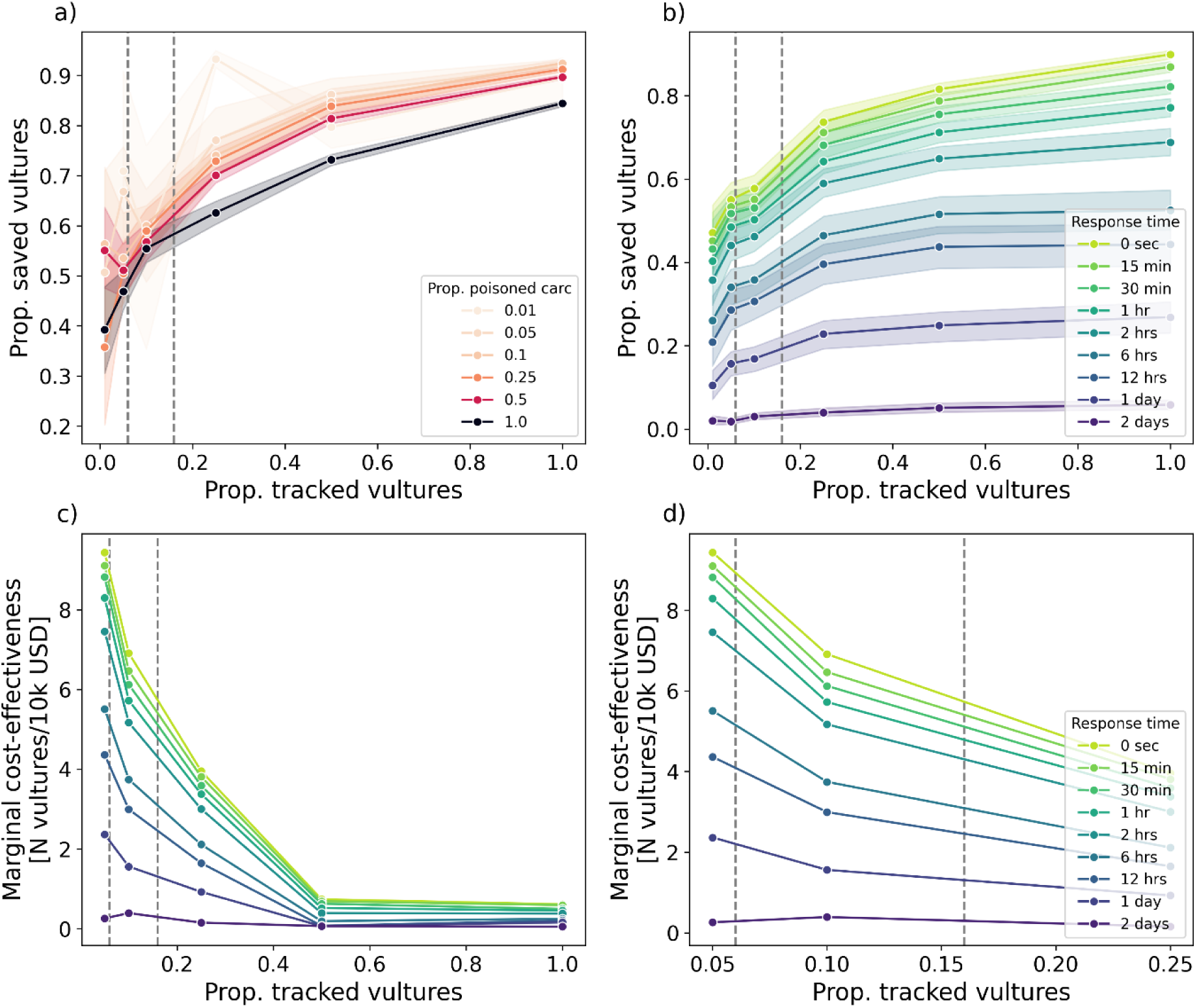
Chains of vultures. (a) Proportion of saved vultures (those arriving at the poisoned carcass after the first tracked individual) under varying proportions of tracked vultures and poisoned carcasses. (b) Proportion of saved vultures with increasing tracked vultures and response times (time after the first tracked individual arrives at the carcass), across all poisoned carcass proportions. (c) Marginal cost-effectiveness showing additional vultures saved per 10,000 USD spent, with increasing tracked vultures and response times, across all poisoned carcass proportions. (d) Zoom-in from panel c for tracked vultures between 0.05 and 0.25. Dashed lines indicate the proportion of tracked vultures in our study area at the beginning (0.06) and currently (0.16).

The total estimated costs for vulture tracking were between 14,094 and 1,109,819 USD for 5 – 496 vultures equivalent to the proportion of tracked vultures of 0.01 – 1.00, respectively (see Suppl. mat.). Marginal cost-effectiveness decreased sharply with increasing proportion of tracked vultures and longer response times (Fig. 3c, d). This suggests that the additional benefit (in terms of vultures saved) per extra dollar spent diminishes significantly as more vultures in the population are tracked. Results also indicate that the delay in response time further reduces the effectiveness of saving vultures through tracking them. This indicates diminishing returns, whereby the cost of saving additional vultures outweighs the benefits, especially as more vultures are tracked and response times increase. An optimal proportion of tracked vultures is thus 5% for across most response time scenarios, with the only exception being the response time of 2 days. In this scenario, the optimal proportion of tracked vultures is 10%.

## Discussion

### The impact of social foraging on poisoning risk

Social foraging strategies resulted in a higher proportion of the vulture population feeding at carcasses and, consequently, a higher proportion of poisoned individuals, compared to the nonsocial strategy. This strongly suggests that social foraging greatly increases poisoning risk in vultures. These findings align with those of a previous study, which reported that vultures employing social foraging strategies were more likely to feed at carcasses than those using nonsocial strategies (Cortés-Avizanda et al., 2014). Here, we take it a step further by demonstrating, for the first time, the consequences of these strategies in terms of mortality risk associated with poisoned carcasses.

The two social submodels, local enhancement and chains of vultures, yielded similar results. This similarity likely occurred because vulture chains rarely form at the vulture and carcass densities used in this study, making nonsocial and local enhancement strategies more prevalent. Indeed, the absence of vulture chain formation is expected in areas with a high density of resources and a small to moderate density of foraging vultures (Curk et al., 2025). Our empirical estimate for the proportion of vultures feeding in a day was closest to the nonsocial strategy, implying that vultures in our study area mainly use nonsocial strategies. However, our empirical estimate for the proportion of poisoned vultures suggests that vultures in our study area use both social and nonsocial strategies. This is consistent with a recent study showing that white-backed vultures in ENP can employ different foraging strategies depending on the density of vulture and carcasses in the environment (Curk et al., 2025). This finding also suggests that vulture species using social foraging strategies are at greatest risk of poisoning, potentially affecting population dynamics and persistence.

### Return on investment from sentinel vultures

Vulture tracking significantly contributed to the effective detection of poisoning events, with the number of saved vultures increasing non-linearly as the number of tracked individuals increases. The highest return on the investment, i.e. vultures saved per unit cost, occurred when the percentage of tracked vultures increased from 1% to just 5% of the whole population. Beyond this point, the cost of tracking additional vultures outweighed the benefits. These findings are crucial for informing vulture and scavenger conservation policies and practices. They suggest that, to reduce poisoning-related mortality, it is not necessary to GPS track a large proportion of the local population, but rather a small subset of individuals which would act as sentinels for all the others. The key then lies in promptly responding to the signal provided by the sentinels, which will further contribute to reducing mortality by rapidly decontaminating the site from poison.

In the case of the ENP, under the proportion of vultures tracked (6 – 16%), we can save up to 65% of vultures from poisoning. If intervention occurs within 1 hour, more than half of the vultures can be saved, while interventions within 6 hours can save about 40%. Similarly, with the most cost-effective tracking rate of 5%, we can prevent up to 60% of vulture mortalities, with up to 50% saved if the response time is within 1 hour and about 35% if within 6 hours. The e-obs tags we used allow data uploads up to three times a day. For instance, if data uploads are set for 14:00 and a vulture feeds on a poisoned carcass at 13:30, we would receive the data 30 minutes later. This delay can range from a few minutes to several hours, assuming the data upload times are set within the window where the study species is the most active and expected to be feeding. Once the data are received, they must be analyzed to identify vulture mortality, and the information communicated to the field team. The response timing largely depends on how quickly and accurately we can identify dead vultures and how fast rangers can reach the target location and decontaminate the poisoned carcass. This is influenced by factors such as the location’s accessibility, distance from roads, and the availability of personnel, and can take several hours.

If the parameters are calculated for the specific species and study area from the tracking data, field observations or other sources, our model is applicable to other study systems. However, the optimal proportion of tracked vultures, in terms of cost-effectiveness, depends on the density of vultures and carcasses in the area, as well as the costs of tracking. These costs vary based on the country of the base institution, the fieldwork area, and the resolution or amount of collected data.

### Poisoning risk and early warning systems

Many poisoning cases remain undetected and unreported, making it difficult to determine the frequency of poisoning in our study area. With an estimated population of 496 vultures (as in our study; see Suppl. mat.), a few poisoning events could potentially eradicate the entire local population due to the social foraging nature of vultures. A population viability study in Kruger National Park, South Africa, predicted drastic population declines and potential extinction of white-backed vultures in the coming decades (Murn & Botha, 2017). Poisoning of adults has a disproportionate impact on population persistence in such long-lived species like vultures (Sanz-Aguilar et al., 2017).

Establishing a system for early detection of poisoning, where sentinel vultures facilitate poisoning detection, especially in remote areas, could cost-effectively prevent mass mortalities. Current systems used in several African countries for detecting carcasses that could potentially be used also for detecting vulture mortalities are based on GPS data with 15 min sampling rate.

Within these systems, through the Movebank (https://www.movebank.org) platform, data are fed to MoveApps (https://www.moveapps.org/) where vulture feeding sites are identified. Here additional steps would be needed to mark potential vulture mortalities based on activity at carcasses inferred from GPS and ACC data. Finally, the potential poisoning locations are shared with rangers through live systems (such as EarthRanger: https://www.earthranger.com/) available on their phone or computer. Rangers can then visit the identified sites. The entire process could take up to 30 minutes, excluding the time needed for rangers to reach the specific location.

To reduce response times further, real-time tags with on-board automatic mortality detection inferred from ACC data would be highly beneficial. Since mortality is expected to be a distinct behavioural state, a potential approach would be the use of anomaly detection. Different algorithms can be trained to detect when data differ from a “normal” state (Ruff et al., 2021). A “simple” model like a one-class Support Vector Machine (Schölkopf et al., 2001) could potentially run on the logger’s hardware. Such a system would also help detect poaching and other causes of vulture mortality, such as drowning and persecution.

Several early warning systems have been suggested to prevent human-wildlife conflict and wildlife crime, including automated detection of elephant rumbles using acoustic data (Zeppelzauer & Stoeger, 2015), GPS tracking of lions to alert communities (Weise et al., 2019), using non-targeted sentinel animals to detect poachers (de Knegt et al., 2021), and using Griffon vultures (*Gyps fulvus*) to locate mortality of wild animals and potentially poaching and poisoning (Stoynov, Peshev & Grozdanov, 2018).

In addition to tracking animals and establishing an alert system for early detection of vulture poisoning, efforts should be made to prevent pesticide accessibility and misuse. Currently, agricultural pesticides are still easily available in most African countries, and are used off-label to poison carnivores. Measures should be taken to ban or control the distribution and use of pesticides, enforce penalties for misuse, strengthen law enforcement, increase education, international collaboration and conservation funds (Ogada, 2014; Curk et al., 2024). Additionally, establishing vulture safe zones where targeted conservation measures are implemented could be beneficial, following successful examples in Asia (Safford et al., 2019).

## Conclusions

We show that social foraging predisposes vultures to severe poisoning risks, but we also provide evidence that just a few sentinel birds in a population can save many other birds from poisoning. On a more practical level, we quantify the return on investment of sentinel vulture tracking in relation to the time it takes to decontaminate a poisoned carcass. The findings provide clear guidance for conservation practitioners to mobilize resources and design poison mitigation strategies based on sentinel vultures. At the local scale, mitigating poisoning risk with limited resources will entail a hard trade-off between increasing the number of sentinels versus increasing the number and locations of rangers to rapidly access poisoned carcasses. In an optimal scenario, practitioners should target the tracking of 5% of vultures in the local population, and aim to reduce the time to access poisoned carcasses within 2 hours. This would prevent about 45% vulture mortalities and would represent the most optimal return on investment. Finally, under a chronic lack of conservation resources, particularly in the Global South, providing clear numbers in terms of costs and the ecological benefits will also help securing funds for conservation. For example, an estimation that a budget of 58,728 USD is needed for tracking of 5% of vulture population (25 individuals) will help reduce poisoning mortality by 45% when response time is within 2 hours may represent an extremely powerful and attractive opportunity for potential funders to support. This evidence carries extremely timely and practical implications for designing conservation efforts on the ground to mitigate the consequences of poisoning.

