## Supplementary material for "Using animal tracking for early detection of mass poisoning events"

**Model and parameter description – adapted from Curk et al. (2025)**

### MODEL DESCRIPTION

To describe the agent-based model used in this study, we followed the Overview, Design concepts, and Details (ODD) protocol (Grimm et al. 2006, 2010, 2020, Railsback & Grimm 2019). Table S1 contains the abbreviations used throughout the text.

#### Overview

*Purpose and Patterns* – This model aims to evaluate (1) how different foraging strategies i.e. nonsocial, local enhancement and chains of vultures affect mass poisoning risk in vultures and (2) whether vulture GPS-tracking can be used for early detection of mass poisoning.

For the first objective (1), we incompared three submodels depicting distinct foraging strategies: "nonsocial," where vultures rely solely on personal information to locate carcasses; "local enhancement," where vultures use both personal and social cues to find carcasses by spotting either an unoccupied carcass or other vultures feeding or landing at the carcass; and "chains of vultures", which expands on nonsocial and local enhancement by allowing vultures to follow each other in sequences, potentially forming long chains. We run the submodels several times, each time with different percentage of poisoned carcasses (1%, 5%, 10%, 25%, 50% or 100%). From the output of the there submodels, we derived the proportion of individuals feeding within the first day by calculating the number of vultures feeding out of the number of all vultures included in a simulation. From the output of each submodel, we also extract the proportion of poisoned vultures at the end of 3-day simulation, calculated as the number of poisoned vultures out of the total number of vultures in a simulation run.

For the second objective (2), we used only "chains of vultures" submodel since it includes nonsocial as well as social strategies and in the previous study (Curk et al., 2025) we found that vultures in our study area are able to use nonsocial as well as social strategies, depending on the vulture and carcass density in the environment. We ran the "chains of vultures" submodel with different percentage of poisoned carcasses and tagged individuals (1%, 5%, 10%, 25%, 50% or 100%). From the output of the submodel, we calculated the proportion of saved vultures considering various response times: 0 seconds, 15 minutes, 30 minutes, 1 hour, 2 hours, 6 hours, 12 hours, 1 day, and 2 days. A response time of 0 indicates the moment the first tagged vulture reaches the carcass, while a response time of 15 minutes indicates 15 minutes after the first vulture's arrival, and so on. It is assumed that the carcass is decontaminated after the specified response time, and vultures that would have arrived after the response time if the carcass was not decontaminated are regarded as "saved" from poisoning. Therefore, the proportion of saved vultures is calculated as the number of vultures arriving at carcasses after the response time divided by the total number of vultures arriving at the carcasses during the simulation run.

*Entities, State Variables, and Scales* − The model entities include vultures and carcasses. Vultures are characterized by their x and y coordinates and status, which ranges from 0 (foraging) to 4 (feeding) and 5 (poisoning), with intermediate states indicating various approaches to carcasses. Carcasses are identified by coordinates, poisoning (not poisoned, poisoned), and status (unoccupied, detected, occupied).

The model simulates a 152 x 152 km area, representing Etosha National Park (without Etosha Pan) and 10 km belt of communal farmland around the park. We used a 8 x 8 km cell grid which allows us to speed up calculations of distances between vultures and carcasses. The simulation has 10-second time steps and lasts for three days where one foraging day lasts 5 hours. Periodic boundaries were included to prevent edge effects.

*Process Overview and Scheduling* − Vulture status is initially: 0 – foraging. At each time step, when the vulture is foraging, the model notes the vulture's cell and neighboring cells, calculating distances to carcasses and other vultures. Distances to carcasses are prioritized, followed by distances to vultures based on submodel rules. If within detection distance of a carcass (*Do*, *Du*), a vulture (*Dv*), or a follower (*Df*), the vulture changes direction and approaches the target, changing carcass status from 0 – unoccupied to 1 – detected and vulture status to 1 – approaching an unoccupied carcass, 2 – approaching an occupied carcass or individual landing at the carcass, or 3 – following the vulture that is approaching another individual who is directly or indirectly approaching the carcass. The "nonsocial" submodel includes only *Du*, "local enhancement" includes *Du*, *Do*, and *Dv*, and "chains of vultures" includes all detection distances. Vultures are assumed to lose altitude while approaching the carcass from detection distance, so no extra steps for descent are included. It is assumed that vultures lose altitude as they approach the carcass from the detection distance, so no additional steps for descent are included in the model. When the vulture arrives at the carcass, the carcass status updates to 2 – occupied, and the vulture's status changes to 4 – feeding. If the carcass is poisoned, the vulture's status shifts to 5 – poisoning, and it remains at the carcass until the simulation ends (we assume that the vultures do not feed more than once a day). If the carcass is not poisoned, the vulture's status resets to 0 – foraging the next day, and the states can continue to change as previously described. Detailed submodel processes are illustrated in Fig. S1.

Observer processes capture data on vulture IDs, vulture status, whether the vulture is tagged or not, carcass ID the vulture is feeding on or approaching, carcass status, whether the carcass is poisoned or not, and time when the vulture detected the carcass. The data also include indormation about the proportion of tagged individuals and proportion of poisoned carcasses for the specific simulation run. The model outputs data tables at the end of each day, which are used to calculate output variables including, the proportion of individuals feeding, proportion of poisoned vultures and proportion of saved vultures from potential poisoning.

Table S1: State variables and parameters (adapted from Curk et al., 2025)

| Variables/ parameters | Description | Value | Information obtained |
|---|---|---|---|
| *As* | Area size | 152 x 152 km | Etosha National Park with surrounding commercial farmland |
| *Nv* | Number of vultures | 496 | Empirical data and Monadjem, Botha & Murn (2012) |

| | | | |
|---|---|---|---|
| *Nc* | Number of carcasses | 141 | Empirical data, Bellan et al. (2013) |
| *Sd* | Simulation duration | 3 days (5 h per day) | Empirical data |
| *Ts* | Time step | 10 s | Cortés-Avizanda et al. (2014) |
| *Hi* | Initial heading – orientation at the start of simulation | 0 – 360° | Random value drawn from uniform distribution |
| *Hc* | Change in heading per time step | 17° | Empirical data |
| *Fs* | Foraging speed | 15 m/s | Empirical data |
| *Fh* | Flight height | 500 m | Empirical data |
| *Du* | Detection distance to an unoccupied carcass | 300 m | Jackson, Ruxton & Houston (2008), Cortés-Avizanda et al. (2014) |
| *Do* | Detection distance to an occupied carcass | 4,000 m | Jackson, Ruxton & Houston (2008), Cortés-Avizanda et al. (2014) |
| *Dv* | Detection distance to a vulture approaching the carcass | 2,000 m | Empirical data |
| *Df* | Detection distance to the follower | 2,000 m | Empirical data |

### Design Concepts

*Design Concepts* − The *basic principle* is to test whether poisoning risk differs between vultures with different foraging strategies (nonsocial, local enhancement, and chains of vultures) and whether vulture tagging can facilitate with preventing mass poisoning by early detection of poisoning events. Previous modelling studies investigated different foraging strategies of vultures but not in the contex of poisoning.

*Emergence* − The number of vultures aggregating at carcasses and variables, proportion of individuals feeding, proportion of poisoned vultures and proportion of saved vultures, *emerge* from individual behaviours. The model lacks *adaptive behaviour*, *learning*, or *prediction* mechanisms.

Vultures *sense* carcasses and each other within set distances (*Du*, *Do*, *Dv*, *Df*) and adjust their movements accordingly.

*Interactions* occur in the "local enhancement" and "chains of vultures" submodels, where vultures detect others feeding or approaching carcasses. These interactions influence movement decisions and aggregation.

*Stochasticity* − is introduced through random initial positions of vultures and carcasses, as well as initial headings of vultures (*Hi*).

*Observations* − the data are recorded at the end of each day in a simulation, with outputs formatted into tables for metri calculation of the output variables (the proportion of individuals feeding, proportion of poisoned vultures and proportion of saved vultures). We also compare model outputs with empirical data.

There are no *collectives* in the model.

### Details

*Initialization* − Simulations start with random coordinates for carcasses and vultures. Vultures have status foraging and carcasses status unoccupied. Prior to simulation start, the vultures are randomly assigned into tagged or not tagged and carcasses into poisoned or not poisoned, depending on the proportion of tagged individuals and proportion of poisoned carcasses in the specific simulation run.

*Input* – Environmental conditions are not included. The model assumes a static environment.

*Submodels* – We incorporated three submodels representing foraging strategies of vultures (Fig. S1):

"Nonsocial": Vultures switch from foraging (status 0) to approaching unoccupied carcasses (status 1) when within detection distance (*Du*). Upon reaching the carcass, status changes to feeding (status 4) and in cases when the carcass is poisoned, to poisoning (status 5).

"Local enhancement": Vultures first forage (status 0) and transition to approaching unoccupied carcasses (status 1) if within *Du*, or status 2 - approaching occupied carcass or following individual who is approaching the carcass (if the distance to the occupied carcass is less than *Do* or the distance to the vulture that is approaching the carcass is less than *Dv*). Status changes to feeding (status 4) upon arrival and poisoning (status 5) if the carcass is poisoned.

"Chains of vultures": Includes statuses 0, 1, 2, and 4 as in "local enhancement" submodel, plus status 3 - following the vulture that is following another individual. This status initiates indirect carcass approach by following other vultures within *Df*.

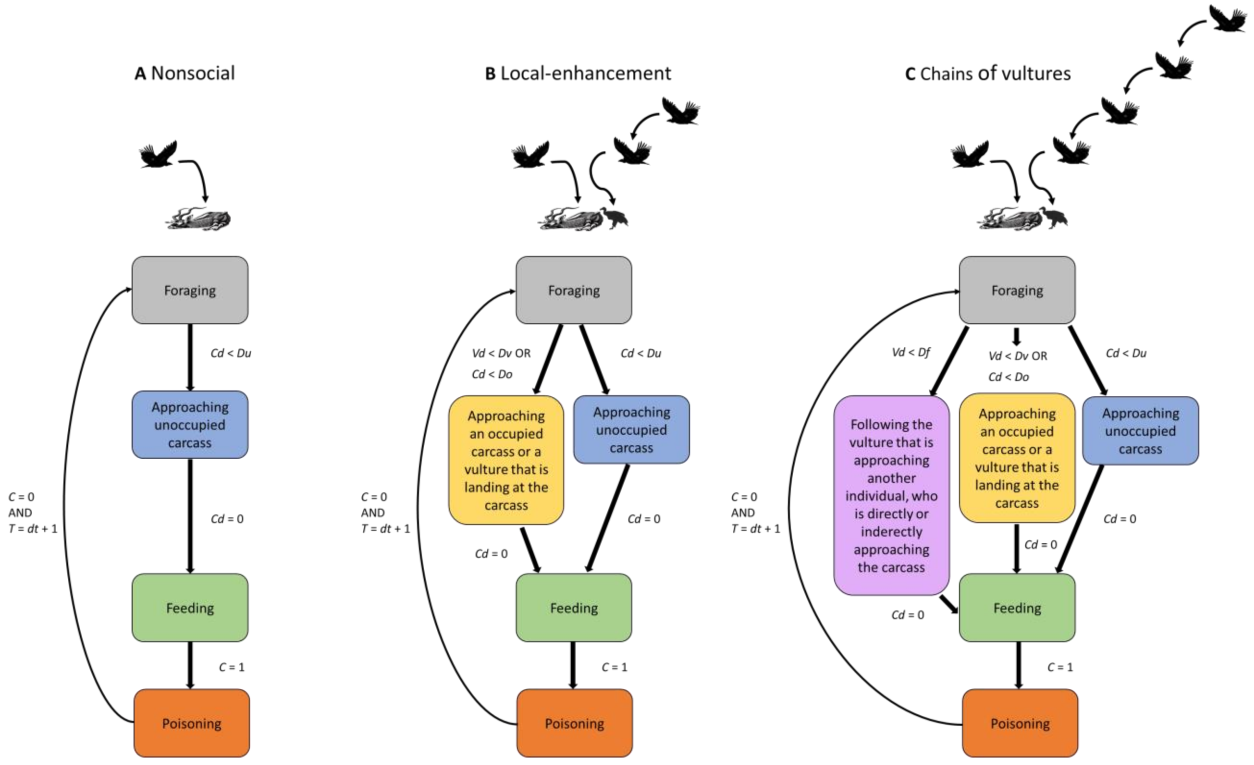


Figure S1: Flowchart of the three submodels adapted from Curk et al. (2025). Vulture status changes depending on the submodel between foraging; approaching unoccupied carcass; approaching occupied carcass or following individual who is landing at the carcass; following the vulture that is approaching another individual who is directly or indirectly approaching the carcass; feeding; and poisoning. *Cd* is distance to the occupied or unoccupied carcass and *Vd* distance to a vulture who is landing at the carcass or a vulture who is approaching another individual who is directly or indirectly approaching the carcass. *C* is carcass status (0 – not poisoned, 1 – poisoned), *T* simulation time and *dt*, current date. *Du* is distance to an unoccupied carcass (300 m), *Do* distance to an occupied carcass (4,000 m), *Dv* distance to a vulture approaching the carcass (2,000 m), and *Df* distance to the follower (2,000 m).

PARAMETER DESCRIPTION

*Number of vultures (Nv)* − We conducted an aerial survey covering 80% of Etosha National Park which identified 84 active nests (egg, chick or an incubating parent present in the nest) of unknown vulture species. Based on this coverage, we estimate that a complete survey (100% coverage) would detect approximately 105 nests within the park. To account for nests on commercial farmland surrounding the park, we expanded the park area in the model by 10 km on each side, resulting in a total survey area of 23,104 km² (152 km x 152 km). Within this extended area, we estimated that there are approximately 139 nests, calculated by proportionally adjusting the number of nests found within the park to the larger survey area.

Based on the estimated population size of white-backed vultures (WBV), lappet-faced vultures (LFV), and white-headed vultures (WHV) in Namibia (Brown, Simmons & Kemper, 2015; Brown & Simmons, 2015; Simmons, 2015; Simmons & Brown, 2015), the proportion of species is as follows: WBVs comprise 87.7% of the population, LFVs comprise 8.8%, and WHVs comprise 3.5%. We did not include hooded vultures since population estimation is only 25 pairs and the nests are usually hidden under the canopy and difficult to be detected from the aircraft. Consequently, we estimated there are 121 WBV nests in the extended survey area for the given season.

For a given year, the vulture population within the extended area was calculated using the following assumptions: Each nest corresponds to a breeding pair, thus there are 242 adult WBVs (121 nests x 2 adults per nest), each breeding pair produces one chick per year, and adults breed annually. Based on our breeding success study of GPS-tagged vultures from 2023 in Etosha (n = 10, unpublished data), 60% of chicks survive until fledging. Therefore, there are approximately 73 first-year birds (121 nests x 60% survival). With an annual survival rate of 91% across all age classes (Monadjem, Botha & Murn 2012), we estimated the number of birds in older age classes as follows: 66 second-year birds (73 first-year birds x 91% survival), 60 third-year birds (66 second-year birds x 91% survival), and 55 fourth-year birds (60 third-year birds x 91% survival). The total WBV population across all age classes in the extended area is thus 496 birds (242 adults + 73 first-year birds + 66 second-year birds + 60 third-year birds + 55 fourth-year birds).

*Number of carcasses (Nc)* − To estimate the total herbivore biomass in the study area, we used the biomass layer from Hempson et al. (2015), which includes 92 wild herbivore species across Africa as well as livestock (cattle, sheep, and goats). The total biomass for the area was calculated to be 194,750,000 kg. The average animal weight, including key species in Etosha and livestock, is approximately 761 kg. By dividing the total biomass by this average weight, we estimated that there are around 255,913 animals in the study area.

Using the mortality index based on Bellan et al. (2013), which accounts for a mean mortality rate of 0.1% (considering scenarios with and without anthrax), the daily mortality for wild herbivores was calculated to be 256 animals. Here we assumed the same mortality index for all herbivores. For livestock, the annual mortality rate based on data from 2015 (Santangeli et al., 2016) is 3.8%. This translates to an annual mortality of approximately 9,725 animals. When averaged over the year, the daily mortality rate for livestock is around 27 animals. To obtain a comprehensive daily mortality rate that accounts for both wild herbivores and livestock, we averaged the two daily mortality rates, resulting in an estimated mean daily mortality of 141 animals, which we used as a number of carcasses in our model. Note that in the model we assumed that after the initial set of carcasses, new carcasses do not appear during the three-day simulation period.

*Simulation duration (Sd)* − The duration of a simulation is three days, where each day corresponds to 5 foraging hours. The foraging length of 5 hours was used because the highest speeds of the GPS tracked vultures were observed between 9:00 and 14:00 GPS time, representing a 5-hour period (n = 26,822,948; see Fig S3 in Curk et al., 2025). We used the length of three days, since we observed in the field (personal observation) that most carcasses are eaten within the three days, considering the carcass size of 761 kg as used in the model.

*Change in heading (Hc)* refers to the alteration in the direction of movement between consecutive time steps, considering only flight locations with speeds greater than 2 m/s, and excluding those within 4,000 meters of the carcass. The change in heading per time step was calculated using the R package "move" (Kranstauber et al. 2022). On average, the change in heading was 16.6 degrees (± 0.03). In the model, for each foraging step, a random change in heading was included, drawn from a normal distribution with a standard deviation of 17 degrees (16.6), and the calculations were performed using radians.

*Foraging speed (Fs)* represents the average speed, considering only flight locations where the speed was greater than 2 m/s before reaching the carcass location on that day, and excluding those within 4,000 meters of the carcass (mean ± SE: 14.5 ± 0.004, n = 1,475,405). Speed calculations were performed using the R package "move" (Kranstauber et al. 2022).

*Flight height (Fs)* is the average foraging altitude, considering only flight locations where the speed exceeded 2 m/s before reaching the carcass location on that day and excluding those within 4,000 meters of the carcass (mean ± SE: 529.2 ± 0.3, n = 1,473,737). Flight height was determined by subtracting the height from the DEM (Digital Elevation Model) raster with 30m spatial resolution (ASTGTMV003) from the height above the ellipsoid measured by the GPS tag.

*Detection distances (Du, Do, Dv, Df)* – Based on Jackson, Ruxton & Houston (2008) and Cortes-Avizanda et al. (2014), we used 300 meters as the horizontal detection distance for unoccupied carcasses (*Du*) and 4,000 meters for occupied carcasses (*Do*). The detection distances for a vulture approaching the carcass (*Dv*) and for the follower (*Df*) were inferred from GPS trajectories and carcass locations. Data were used when multiple individuals approached the carcass simultaneously, considering only the first day the carcass was detected. The detection distance to an approaching vulture was defined as the distance from the point where the individual's distance to the carcass started decreasing sharply (see Fig. S2 in Curk et al., 2025). The average distance between the follower and the leader was 2,240 meters (n=16). However, it is possible these individuals were following another non-tagged vulture approaching the carcass. For simplicity, we used a rounded value of 2,000 meters for *Df* in the model.

Note that the empirical values *Sd*, *Fh*, *Fs*, *Hc* and detection distances *Dv* and *Df* were calculated using R version 4.3.0. For simplification, we used rounded values in the model.

### SCENARIOS

First, to examine how various foraging strategies influence poisoning risk in vultures, we ran the three submodels multiple times, each with a different percentage of poisoned carcasses (1%, 5%, 10%, 25%, 50%, or 100%). We conducted 10 repetitions for each percentage, resulting in 180 simulations (3 submodels x 6 poisoned carcass scenarios x 10 repetitions). Second, to evaluate the potential of GPS-tracking for early poisoning detection, we ran the chains of vultures submodel with varying percentages of poisoned carcasses and tagged individuals (1%, 5%, 10%, 25%, 50%, or 100%). This also included 10 repetitions for each scenario, leading to 360 simulations (1 submodel x 6 poisoned carcass scenarios x 6 tagged vulture scenarios x 10 repetitions).

**Costs of vulture tracking**

Objective of costed intervention: This study describes the costs of tracking white-backed vultures (*Gyps africanus*) in Etosha National Park (Namibia) and surrounding communal farmland where poisoning risk is high. GPS-tracking is used for early detection of vulture poisoning events. The project is not yet established and the purpose of the study is to estimate how many vultures can be saved with tracking different proportion of vulture population and evaluate cost-effectiveness of such intervention.

Methodology of costed intervention: The method applied is trapping and equipping vultures with solar units in the field tracking them for one year.

Context of costed intervention: Population of white-backed vultures in the study area was estimated from aerial-survey reporting active vulture nests (see “Parameter description” above).

Intervention scale: Etosha National Park

Duration of intervention so far (years): 1

Was the objective achieved? Not yet

Categories included in costs (further breakdown below): Labor, capital assets, consumables

Describe discounting or inflation correction if applicable: N/A

Organizational level of cost data: Intervention level costs

Total cost of intervention: 5290 USD per vulture, 2023 values

Table S2: Costs of vulture tracking per solar unit and per one individual for one year

| Cost Category | Description | Unit cost | Units | Fixed/Variable | Currency | Date |
|---|---|---|---|---|---|---|
| consumable | Bird solar GPRS/UMTS 42g (e-obs) | 1925 | N/A | variable | USD | 2023 |
| consumable | cellular network access and data transmission via SMS and GPRS/UMTS | 256 | 1 year | variable | USD | 2023 |
| consumable | base station + accessories | 2783 | N/A | variable | USD | 2023 |
| consumable | harness material (teflon) | 20 | 1.5 meter | variable | USD | 2023 |
| labor | 2 field assistants | 293 | 1 day | variable | USD | 2023 |
| consumable | fuel | 13 | 12 liter/100km | variable | USD | 2023 |

Note that the costs of the bird solar unit, network access and data transmission, and harness material increase linearly with the number of tagged vultures, while base station costs remain constant regardless of the number of vultures tagged. The costs of fieldwork (field assistants and fuel) do not increase linearly. We assume that 10 vultures are on average captured and equipped with solar units per working day using cannon-net system. This can also mean that two days are unsuccessful and on the third day, 30 vultures are captured. Preparation, driving and setting-up the trap also takes time thus we assume that one working day is needed for 1 vulture as well as for 5 vultures, 3 days for 25 vultures, 5 days for 50 vultures, 13 days for 124 vultures, 25 days for 248 vultures and 50 days for 496 vultures. We assume that on average 100 km are covered in the field by car per working day.

### Sensitivity analysis

We conducted a local sensitivity analysis by varying one parameter at a time to evaluate how changes in parameter values affect model outputs (Railsback & Grimm, 2019). We utilized the chains of vultures model, which incorporates all relevant parameters, with a simulation duration of 5 hours representing one foraging day. The proportion of tagged vultures and poisoned carcasses was set to 0.5. We assessed the sensitivity of parameters *Nv*, *Nc*, *Hc*, *Fs*, *Du*, *Do*, *Dv*, and *Df*. Each parameter was adjusted to ± 5% of its reference value while the other parameters remained at their reference levels (Table 1). Additionally, we ran the submodel with all variables set to their reference values. This resulted in 170 simulation runs ((8 parameters x 2 sensitivity levels + 1 reference level) x 10 repetitions). From the model outputs, we calculated three metrics: the proportion of vultures feeding, the proportion of poisoned vultures, and the proportion of saved vultures. For details how sensitivities were calculated, see Curk et al. (2025).

The local sensitivity analysis revealed that the output variables from the model were similarly influenced by changes in parameter values (Table S3). However, detection distances *Df*, *Dv*, and *Do* had the most significant impact on the output variables (proportion of saved vultures and proportion of poisoned vultures), with differences greater than 0.5 between sensitivity plus and sensitivity minus. Thus, future studies should take special care to provide precise estimations of detection distances.

Table S3: Sensitivity plus and sensitivity minus for each parameter (*Nv* - number of vultures, *Nc* - number of carcasses, *Hc* - change in heading per time step, *Fs* - foraging speed, *Du* - detection distance to an unoccupied carcass, *Do* - detection distance to an occupied carcass, *Dv* - detection distance to a vulture approaching the carcass, *Df* - detection distance to the follower) and output variable (proportion of vultures feeding, proportion of poisoned vultures, proportion of saved vultures).

| Parameter | Output variable | Sensitivity plus | Sensitivity minus |
|---|---|---|---|
| *Nv* | Prop. feeding | 1.05 | 1.09 |
| *Nv* | Prop. poisoned | 0.57 | 0.81 |
| *Nv* | Prop. saved | 0.46 | 0.38 |
| *Nc* | Prop. feeding | 0.05 | 0.01 |
| *Nc* | Prop. poisoned | -0.21 | 0.02 |
| *Nc* | Prop. saved | -0.29 | 0.1 |
| *Hc* | Prop. feeding | -0.06 | -0.03 |
| *Hc* | Prop. poisoned | -0.36 | 0.04 |
| *Hc* | Prop. saved | -0.23 | -0.4 |
| *Fs* | Prop. feeding | 0.04 | 0.12 |
| *Fs* | Prop. poisoned | 0.49 | 0.01 |
| *Fs* | Prop. saved | -0.09 | -0.13 |
| *Du* | Prop. feeding | -0.03 | -0.01 |
| *Du* | Prop. poisoned | 0.03 | 0.31 |
| *Du* | Prop. saved | 0 | 0.2 |
| *Do* | Prop. feeding | 0.04 | 0.16 |
| *Do* | Prop. poisoned | 0.19 | 0.15 |
| *Do* | Prop. saved | -0.11 | 0.54 |
| *Dv* | Prop. feeding | 0.03 | 0 |
| *Dv* | Prop. poisoned | -0.33 | 0.35 |
| *Dv* | Prop. saved | -0.6 | 0.8 |
| *Df* | Prop. feeding | -0.01 | -0.02 |

| | | | |
|---|---|---|---|
| *Df* | Prop. poisoned | -0.44 | 0.25 |
| *Df* | Prop. saved | -0.45 | 1.01 |
